# Interactive effects of temperature acclimation and dietary fatty acids on metabolic rate and body composition of zebra finches (*Taeniopygia guttata*)

**DOI:** 10.1101/2024.07.30.604977

**Authors:** Michael J. Campbell, Gabriela F. Mastromonaco, Gary Burness

## Abstract

Climate change is contributing to geographic range shifts in many bird species, with possible exposure to novel diets. How individuals respond physiologically across chronic time frames to the interacting effects of diet and environmental temperature has been little explored. We acclimated zebra finches to either cool (20-24°C) or thermoneutral (35°C) temperatures over 6-months and provided them with diets enriched in either unsaturated or saturated fatty acids. We measured body mass throughout the study, and basal metabolic rate (BMR) and body composition at 3- and 6-months, respectively. Individuals held in cool conditions and fed a diet enriched with unsaturated fatty acids lost mass initially relative to the other groups, however, effects were reversible, and all individuals had a similar mass at 6- months. Chronic exposure to cool conditions increased BMR and the mass of the pectoral muscle and visceral organs. However, we could detect no long-term effect of diet on any physiological parameter. Our results contrast with those of birds studied over acute time frames, in which diet and temperature interact to determine energy expenditure. Over chronic time frames individuals appear to reach a new steady-state, with long-term physiological responses driven primarily by thermoregulatory responses to environmental temperature.

**Research Highlights:**

- With climate change, birds may encounter novel diets and temperatures
- In zebra finches we show that chronic acclimation to cool temperatures increased energy expenditure and changed body composition.
- Dietary fatty acid content had little long-term impact on the physiological parameters we measured.

## Introduction

Across taxa, the interacting effects of food availability and ambient temperature can impact the behaviour and physiology of individuals. For example, various species of birds have shown range shifts over time, with individuals tracking their preferred food sources and environmental temperatures (e.g., ptarmigans (*Lagopus*; Lagerholm et al. 2017); *Charadiiformes* Maclean et al. 2008). Meanwhile, members of other species may delay seasonal migration and remain as residents while enduring cold winter temperatures, as long as their preferred food remains available (e.g., Newton 2006; Lindén et al. 2011). These responses are, at least in part, the outcome of attempts by individuals to balance nutrient and energy intake rates against levels of energy expenditure (Stalmaster & Gessaman 1984).

When birds respond to changes in ambient temperature, they often adjust their lipid stores (West & Meng 1968; Li et al. 2020). In cool conditions, increased lipid stores allow individuals to retain heat (Houston et al. 1997) and serve as energy reserves (Cooper 2007). However, the type of lipids consumed in the diet can impact the relative ease with which individuals can maintain fat stores over the long-term. For example, unsaturated fatty acids are mobilized more rapidly than saturated fatty acids (Mustonen et al. 2009). As a result, birds on an unsaturated fatty acid-rich diet may to need to consume food at higher rates or risk diminishing their lipid stores, despite the benefits of being able to rapidly mobilise fuels during times of increased energy expenditure (Guglielmo 2010; Zielinski & Pratt 2017). However, maintaining larger fat stores for long periods of time increases the risk of oxidative stress due to unsaturated fatty acids being more prone to lipid peroxidation (Zielinski & Pratt 2017 & Su et al. 2019).

Although birds use their lipid stores as a source of insulation during short-term daily cycles (Houston et al. 1997; Dubois et al. 2016), long-term temperature acclimation typically involves adjustments in heat production via adjustments in basal metabolic rate (BMR; McKechnie 2008; Swanson 2010) and the size of visceral organs (Liknes & Swanson 2011; Zheng et al. 2014). BMR can be influenced by cell membrane fatty acid composition (Hulbert et al. 2005), which in turn may be affected by a bird’s diet (Guglielmo et al. 2002; Hulbert et al. 2005; McCue et al. 2009; Carter et al. 2019). For example, diets rich in unsaturated fatty acids have an elevated ratio of double bonds to the number of fatty acid chains (i.e., unsaturation index; UIN; Pamplona 2008). By increasing an individual’s UIN, membrane leakage increases which can lower the efficiency of cellular processes and result in an increased release of heat (Roussel 2020). As a result, birds living under cool conditions may preferentially consume diets rich in unsaturated fatty acids (McWilliams et al. 2002) because these diets allow for greater passive heat production (McGuire et al. 2013).

To date, a handful of studies in birds have explored interactive effects of temperature and dietary fatty acids on avian physiology. For example, at low temperatures great tits (*Parus major*) have high BMR when fed a diet rich in saturated fatty acids but not when they were fed diets rich in unsaturated fatty acids (Andersson et al 2018). Meanwhile, at warm temperatures, the reverse pattern occurred, with individuals fed an unsaturated fatty acid rich diet having the highest BMR. Given that individuals were housed at their respective temperature for ∼1 weeks before being measured, the interactive effects of diet and temperature on metabolism can be rapid (Andersson et al 2018).

Temperature acclimation can occur within a relatively short amount of time (1-3 weeks; McKechnie 2008; Dubois et al. 2016; Thompson et al. 2016). In contrast, responding to changes in dietary fat composition may take months before birds fully acclimate their tissues to reflect the diet being consumed (Carter et al. 2019). Despite the large difference in time required to fully acclimate to temperature and diet there have been no studies, to our knowledge, that have examined how individuals respond to both factors over a chronic time frame. In the current study, we explored the long-term interactive effects of temperature acclimation and dietary fatty acid content on metabolic rate and body composition in captive zebra finches (*Taeniopygia guttata*). Zebra finches are a well-characterized, non-migratory granivore, native to Australia and Indonesia. Over the course of 6-months we exposed individuals to either cool or thermoneutral ambient temperatures, while supplementing their diets with either saturated or unsaturated fatty acids. We found that dietary fatty acids had relatively little impact on metabolic rate at 3-months, nor body composition after 6-months, with most effects due simply to acclimation temperature.

## Methods and Methods

### Animal care

All experiments involving the use of animals were reviewed and approved by the Trent University Animal Care Committee (Animal Use Protocol # 26447). All experiments were performed in accordance with Trent University and the Canadian Council on Animal Care guidelines on the care and use of laboratory animals.

Zebra finches were from of colony housed in the Trent University Animal Care Facility. Experimental individuals were housed in one of 16 single-sex flight cages (45 cm × 45 cm × 90 cm), with 4-6 individuals per cage. Cages were within one of two environmental chambers set to 28°C, and maintained on a 12L:12D photoperiod (08:00-20:00; at 0700 lights turned on at 30% power before ramping up to 100% at 0800, and at 1900 decreased to 30% power before turning off completely at 20:00). All individuals were provided with *ad libitum* grit, water and “Canary/Finch Food” seed mix (containing plain canary seed, canola, nyjer, flax, and red villet; Essex Topcrop Sales Limited). All birds received cooked egg daily and lettuce twice per week, but approximately two months into the primary experiment we reduced egg provisioning to every second day.

### Pilot study

We initially conducted a pilot to ensure zebra finches would accept adjustments to their diet and/or environmental temperature. We weighed each individual (n = 96), collected a 100ul blood sample from the brachial vein (as part of another study), and began supplementing the standard seed diet with either sunflower oil (Unsaturated fatty-acid diet [UFA]: PC Organics Organic Sunflower Oil) or coconut oil (Saturated Fatty acid diet [SFA]: Nature’s Way Premium Coconut Oil) at a seed-to-oil ratio of 5:1 (w:w), following Andersson et al. (2018). To compensate for the higher level of vitamin E found in sunflower oil, we added α-tocopherol (Sigma Aldrich (±) α-tocopherol, T3251-25G) to the coconut oil at 50 mg/100 g oil (Shahidi 2006; Andersson et al. 2018). On the fourth day of exposure to the new diet (Day 4), we began adjusting the temperature of the environmental chambers by 3 °C/day, from 28°C to either a cool temperature (18°C) or one within thermoneutral zone (35 °C; Calder 1964; Wojciechowski et al. 2020). This 2 x 2 experimental design resulted in 4 treatment groups. We stopped the pilot after 12 days, because 3 birds died (one on day 3, prior to adjustment of temperature, and 2 birds on day 12); 8 other individuals started to display signs of sickness/discomfort (e.g., rufled feathers and squinting eyes). We returned all birds to the standard seed diet (i.e., Canary/Finch Food” seed mix), and over the span of two days we returned the temperature of the environmental chambers back to 28 °C. Individuals were kept on the standard diet and housed at 28°C for 14 days, at which point we began the primary experiment.

### Experimental diet and temperature manipulation

All birds included in the primary experiment (n = 88) had also experienced the pilot. At the start of the primary experiment (day 0), we adjusted the temperatures of each environmental chamber by 4 °C/day from 28°C to either 20 °C (cool) or 35 °C (Thermoneutrality). The cool temperature is similar to average maximum temperatures wild zebra finches experience in winter (Zann et al. 1995). On day 14, we began supplementing the standard seed by mix with sunflower oil (Unsaturated fatty-acid diet [UFA]) or coconut oil (Saturated Fatty acid diet [SFA]) as we did in the pilot, but this time at a seed-to-oil ratio of 11:1 (w:w), following McCue et al. (2009). As in the pilot, we added α-tocopherol to the coconut oil at 50 mg/100 g oil (Shahidi 2006; Andersson et al. 2018). This resulted in 4 treatment groups: UFA-enriched diet x Cool temperature (UFA-cool, n = 10 females, 8 males); SFA-enriched diet x Cool temperature (SFA-cool, n = 12 females, 11 males); UFA-enriched diet x Thermoneutral temperature (UFA-TNZ, n = 11 females, 12 males); SFA-enriched diet x Thermoneutral temperature (SFA-TNZ; n = 12 females, 12 males).

Approximately 1 month after beginning experimental treatment (day 34), 8 individuals in the UFA-cool temperature treatment began to display roused feathers and squinted eyes; we removed these individuals from the experiment, and we raised the temperature of the environmental chamber housing the cool treatment birds (UFA-cool and SFA-cool) from 20°C to 24 °C. No further adjustments to temperature or diet were made. Each treatment group had a replicate cage (with 4-6 individuals per cage). However, after 2 months of acclimation we merged the two UFA-cool male cages into a single cage to ensure a similar density of individuals among treatments.

### Blood draws and body mass

We weighed all individuals (± 0.001 g, Sartorius LE 2202) and collected a 100ul blood samples prior to the pilot (T_Initial_, 28-30 days prior the start of the primary experiment) and again every 4-6 weeks over the 6-month primary experiment. We report initial, 1-month, and final (6- month) body mass in this manuscript; data from the blood samples are reported elsewhere (Campbell 2024).

### Basal metabolic rate (BMR)

Between 3-5 months (day 85-137) after the start of the primary experiment we measured the basal metabolic rate (BMR) of a subset of individuals using flow through respirometry, following Burness et al. (2010). As an index of BMR we measured an individual’s post-absorptive, overnight oxygen consumption rate, at thermoneutrality. Individuals were fasted for 1.5 hours prior to being placed into one of three 1 L plexiglass chambers, within a temperature-controlled cabinet set to 35 °C (within the thermoneutral zone: Calder 1964; Wojciechowski et al. 2020). Pre- and post-respirometry body masses of each bird were recorded for each trial. Rates of O_2_ consumption and CO_2_ production and were measured overnight starting between 19:00 and 21:30.

Air was scrubbed of water vapor and CO_2_ using indicating Drierite, soda lime, and Ascarite prior to entering the metabolic chamber. Air flow to the 3 chambers and a baseline (containing no chamber) was set to 500 mL/min and was directed by a multiplexor (TR-RM, Sable Systems, Las Vega, NV, USA) and regulated using mass flow controllers (Sierra Instruments Inc). Air leaving each chamber was subsampled and scrubbed of water vapor prior to reaching the CO_2_ analyzer (CA-10, Sable Systems, Las Vega, NV, USA), and of CO_2_ prior to entering the O_2_ analyzer (FC-10, Sable systems, Las Vegas, NV, USA). The system was calibrated before every run using >99% N_2_ gas, 1.00% CO_2_ gas, and room air scrubbed of water vapor and CO_2_ (20.95% O_2_). The accuracy (± 3%) of the respirometry system was verified every second run via burning a known amount of methanol.

During an overnight respirometry trial, each chamber was sampled for 20-minutes and then the multiplexor switched to record the baseline for 20 minutes, thereby allowing for the analyzers to be flushed and to account for any drift. Concentrations of O_2_ and CO_2_ were recorded every 5 seconds, with the BMR for each bird estimated as the average of the lowest consecutive 50 samples (4 minutes 10 seconds) through the whole night. We calculated VO_2_ and VCO_2_ using equations 10.1 and 10.7 from Lighton (2008), respectively.

### Proportion of time spent feeding

Birds were group housed, so it was not possible to measure individual food consumption. However, as a proxy for food consumption, we video-recorded individual feeding events. Between 183-203 days after the start of the experiment, we placed a video camera (Activeon, CCA10W, Activeon Inc.; San Diego, CA, USA) in front of each cage and recorded individual movements within the cage for approx. 2-hrs at a time. Cages and times of day during which recordings were made were selected randomly.

We discarded the first 30, 60, or 90 seconds of each video recording (depending on the time required to leave the room once we deployed the camera). Every 30 seconds thereafter we scored whether individuals were at the feeder or not. If a bird was present at the feeder, we examined the 10 seconds before and after the 30-second mark to assess whether the bird was actually feeding or not. All but 7 individuals could be individually identified using band color and plumage. These 7 individuals were spread across three cages (two pairs and a triplet). When we observed one of these 7 individuals feeding at the feeder, we added the feeding event to the cumulative total of feeding events for their cage. For subsequent analysis, we assigned each of the twins or triplet birds the feeding rate equal to the total number of feedings observed for their cage, divided by 2 or 3, respectively.

### Organ and fat mass

To quantify the chronic effects of diet and temperature on an individual’s body composition we euthanized a subset of individuals (UFA-Cool: n = 7; UFA-TNZ: n = 14; SFA-cool: n = 13; SFA-TNZ: n = 14). We weighed each individual and collected a final blood sample (mean = 196.4 days; range: 189-203 days). Individuals were anaesthetized with isoflurane gas and then euthanized via decapitation. Within 5 minutes of euthanasia, the left pectoralis muscle, heart, and liver were dissected and placed in pre-weighed air-tight cryotubes, frozen in liquid nitrogen, and then transferred to -70 °C for storage. The gizzard and small intestine were cut open longitudinally, flushed of debris using tap water, and patted dry with paper towel. Wet mass for right pectoralis, kidneys, gizzard, and small intestines were measured on a pan balance (± 0.001 g, SI-124 Denver Instrument). After weighing, the right pectoralis, kidney, gizzard, and small intestine were returned to the carcass and stored at -20 °C for fat content analysis. Wet mass of the heart and liver was determined as the difference between the mass of the vials containing the heart and liver and the pre-determined mass of the vials (the left pectoralis was not weighed and remained frozen).

Carcasses (minus the left pectoral muscle, heart, and liver, which were saved for a separate study) were placed in ashless filter paper envelopes and freeze-dried to a constant mass for 72 hours. To estimate fat content, dried carcasses were placed into a Soxhlet apparatus containing petroleum ether (40/60) for 8 hours, and then air dried in fume hood for 8 hours (to evaporate any remaining solvent). Fat content was calculated as the difference between pre-and post-fat extraction mass.

### Statistical Methods

Statistical analyses were carried out using factoextra (v.1.0.7, Kassambara and Mundt 2020), lme4 (v.1.1.31, Bates et al. 2015), AICcmodavg (v. 2.3.1, Mazerolle 2020), and MuMIn (v. 1.47.1, Bartoń 2022), in R statistical software v.4.2.2 (R Core Team 2022).

### Information theoretic analyses

To identify which independent factors explained the most variation in physiological traits we used an information theoretic approach (Burnham et al. 2010). Models were evaluated using corrected Akaike Information Criterion (AIC_C_). Models with a ΔAIC_C_ of < 2 were considered to hold strong support (Burnham & Anderson 2004). R^2^ was used with ΔAIC_C_ and evidence ratios to provide information on the goodness of fit for each set of models. When we performed linear mixed models, we report conditional and marginal R_2_ values (accounting for, and not accounting for variation due to the random effect, respectively; Nakagawa & Schielzeth 2013). When we tested for variation in right pectoralis (here after, pectoralis), carcass fat, and mass of visceral organ we instead used linear regressions (rather than mixed models) because the random effect of cage identity explained less than a thousandth of the variation. In the cases where linear regressions were used, we report adjusted R^2^ values. To display the directionality of the trends, 95% confidence intervals (CI) and associated coefficients are reported.

### Body mass, pectoralis mass, organ mass, and fat mass

To explore factors contributing to variation in body mass, we performed linear mixed models, including the random effects of cage, and individual identity nested within cage (Schielzeth & Nakagawa 2013). As independent predictors, models included combinations of temperature (cool or thermoneutral [TNZ]), diet (Unsaturated fatty acids [UFA] or Saturated Fatty acids [SFA]), time (initial, 1-month, and 6-months, as ordinal variables), and sex. Additionally, we examined models including the two and three-way interactions of diet, temperature, and time.

Pectoralis muscle mass was analyzed separately from the visceral organs because it is functionally distinct, contributing largely to shivering thermogenesis and peripheral movement. To analyze variation in the mass of visceral organs (heart, liver, gizzard, small intestine, kidney) we first performed a Principal Component Analysis (PCA) using a correlation matrix to collapse the data into orthogonal axes. Principal component 1 (PC1, n = 42 individuals) explained 55.2% of the variation in organ size and had an eigenvalue of 2.762. All organs loaded positively, with correlations ranging from 0.634 (small intestine) to 0.847 (heart).

To analyse variation in the mass of visceral organs (PC-scores), pectoralis, and carcass fat, we performed linear regressions. We tested for the main effects of diet, temperature, body mass, sex, the two-way interactions of body mass, diet, and temperature, and the two-way interaction of body mass with sex. In the analyses of fat mass or pectoralis mass, prior to inclusion of body mass as a main effect we first subtracted fat mass or pectoralis mass from body mass (depending on the analysis), to avoid part-whole correlations (Christians 1999).

### Basal metabolic rate (BMR)

Six birds had biologically improbable resting respiratory exchange ratios (e.g., negative values or > 1.0). We re-measured the BMR of four of these birds on a separate night and used this second reading in our subsequent analysis; we excluded the other two birds as outliers as we could not re-run them. In the end, we had 71 individuals with reliable estimates of O_2_ consumption. The statistical models for assessing variation in BMR included cage identity and respirometry chamber identity as crossed random effects. Models included the main effects of diet, temperature, body mass (averaged from before and after a respirometry trial), and sex as fixed effects, as well as the two- and three-way interactions between diet, temperature, and mass.

### Proportion of time feeding

For each individual we calculated the proportion of observation periods (every 30 seconds, over ∼2-hours) during which that individual was observed to be feeding. One individual (of 64) had an unusually high feeding frequency (> 4SD of the mean). This individual was omitted from further analysis, as a probable outlier. Data were arcsine square root transformed to improve model assumptions. We tested for variation in proportion of time spent feeding using a linear mixed model including temperature, diet, and their interaction as fixed effects, and cage identity as a random effect.

## Results

### Body mass varied with temperature, diet, and length of acclimation

Body mass was best predicted by the length of time an individual had been in the experiment, as “Time” appeared in each model with strong support (ΔAIC_C_ < 2.0; Table 1). There was also a strong 3-way interaction between temperature, diet, and time (ΔAIC_C_ = 0.21, Table 1). At 1-month, birds in cool conditions and fed a UFA-rich diet (UFA-Cool) lost mass, while those fed a SFA-rich diet (SFA-Cool) retained mass (Fig 1; β = 2.512, P<0.001, Table A1). By 6-months, there was a trend for SFA-cool birds to still be heavier than those in the other treatments, however, this was no longer statistically significant (Fig 1; β = 1.494, P = 0.1139, Table A1). On average, cool-acclimated birds were heavier at 6-months than the birds living at thermoneutrality (β = -1.053, P = 0.038, Table A1).

**Figure 1:**
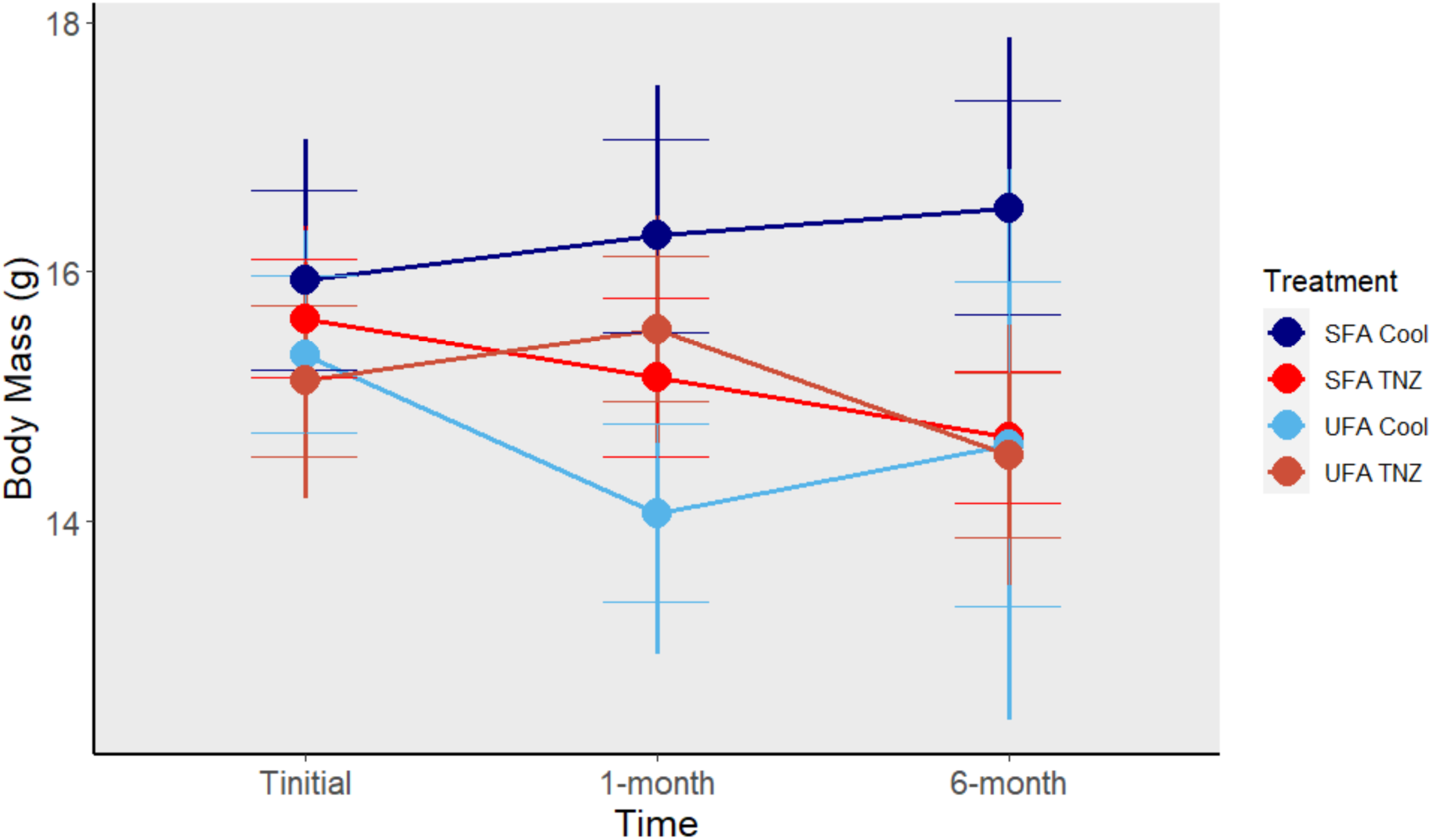
Body mass of zebra finches over the course of a 6-month acclimation to diets enriched in either saturated (SFA) or unsaturated (UFA) fats and held at either cool (24 °C) or thermoneutral (TNZ; 35°C) ambient temperatures. “Initial” sampling period represents the body mass of birds prior to beginning any experimental treatment while other sampling periods reflect birds at approximately 1 and 6 months on treatment. Sexes were pooled. Error bars depict 95% confidence intervals to their full bounds and 80% confidence intervals to within the bounds of the whiskers.

**Table 1:** Model selection results for identifying the factors that explain variation in zebra finch body mass. The table displays only those models with strong support (<2 ΔAIC_C_). Models are ordered by ascending ΔAIC_C_. All models were run as a linear mixed models using cage and individual identity nested within cage as random effects.

| Models | K | AIC<sub>c</sub> | $\Delta AIC_c$ | $W_i$ | Acc $W_i$ | $R^2_m$ | $R^2_c$ | ER |
| --- | --- | --- | --- | --- | --- | --- | --- | --- |
| Time | 6 | 924.74 | 0.00 | 0.17 | 0.17 | 0.014 | 0.733 | 1.00 |
| Temp*Diet*Time | 15 | 924.95 | 0.21 | 0.16 | 0.33 | 0.084 | 0.758 | 1.11 |
| Diet+Time | 7 | 925.26 | 0.52 | 0.13 | 0.46 | 0.040 | 0.733 | 1.30 |
| Time+Sex | 7 | 926.59 | 1.84 | 0.07 | 0.53 | 0.019 | 0.733 | 2.52 |
| Temp+Time | 7 | 926.71 | 1.97 | 0.06 | 0.59 | 0.018 | 0.734 | 2.68 |

Note: K = number of parameters, AIC<sub>c</sub>= corrected Akaike Information Criterion, $\Delta AIC_c$ = difference of focal model AIC<sub>c</sub> compared to the lowest model AIC<sub>c</sub>; $W_i$ = Akaike weight; Acc. $W_i$ = Accumulated weight of models; $R^2_{mar}$ = Marginal R squared; $R^2_{con}$ = Conditional R squared; ER = Evidence ratio; Temp = acclimation temperature (TNZ; 35 °C) or cool (24 °C)); Diet = unsaturated or saturated fatty acid enriched diet; Time = day of experimental exposure (Prior to the experimental treatment, 1-month or 6-month after the start of the experiment). Models with interaction terms also include all internal fixed effects.

### Fat content did not vary with acclimation temperature or diet

At the end of 6-months, an individual’s fat level (carcass fat) was best predicted by ambient temperature, body mass, and sex, with each term appearing in models with strong support (ΔAIC_C_ < 2.0; Table 2). Nonetheless, based on model averaging, temperature did not significantly predict fat mass (Fig 2A; β = 0.578, P = 0.073, Table A2), and fat mass did not vary with body mass nor sex (body mass: β = 0.150, P = 0.325; Sex: β = -0.766, P = 0.710, Table A2). Diet did not appear in any models with strong support.

**Figure 2:**
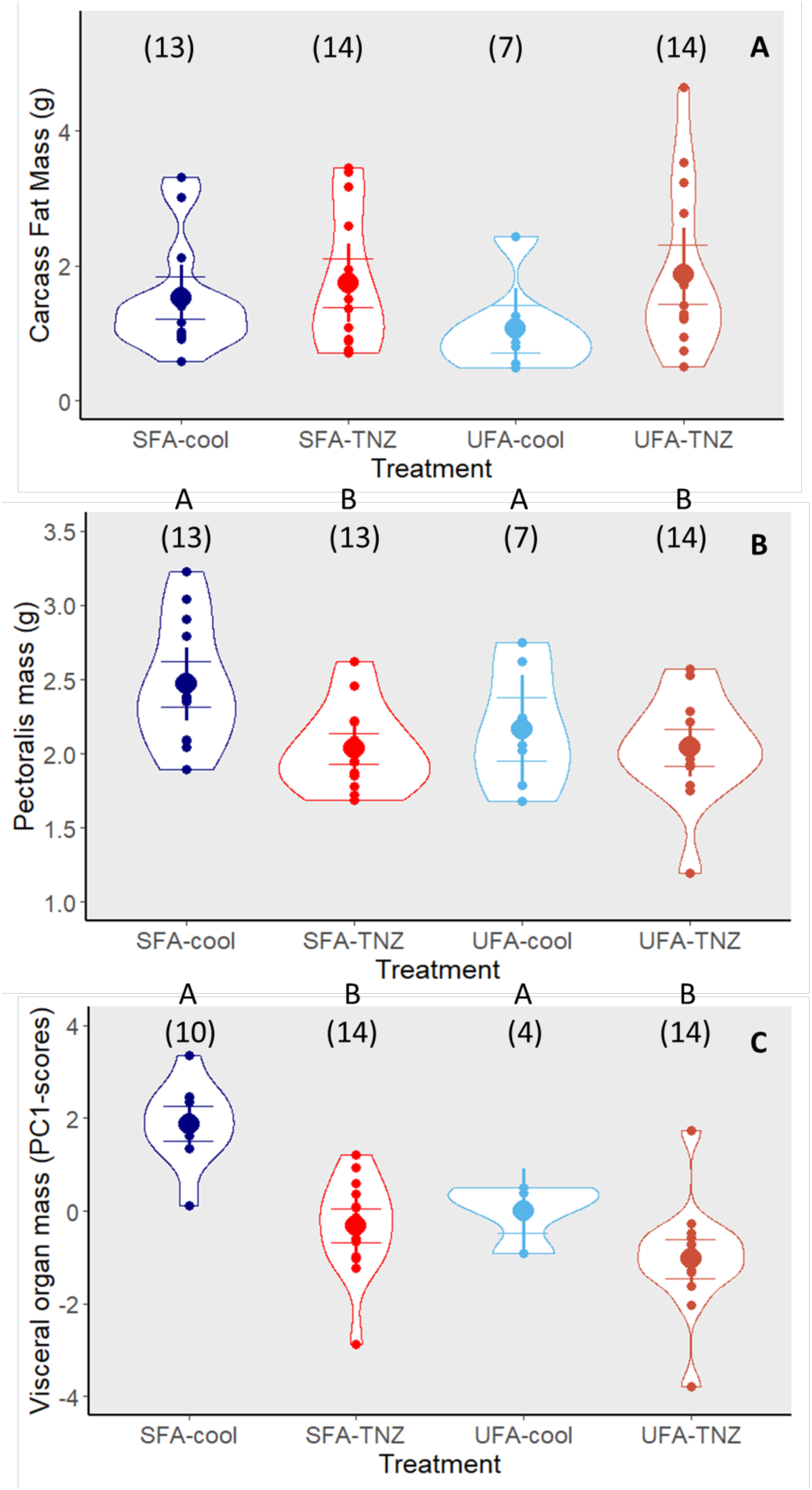
Body composition of zebra finches following a 6-month acclimation to a diet enriched with either saturated (SFA) or unsaturated (UFA) fatty acids, while living at thermoneutral (TNZ; 35 °C) or cool (24 °C) housing temperatures. Mass of (A) Carcass fat; (B) Pectoralis; (C) Visceral organs (PC1 from a Principal Component Analysis, of heart, liver, gizzard, kidney, and small intestine). The large points represent the means, while the error bars represent both the 80% confidence interval (to whiskers) and 95% confidence interval (full range of error bars). Different letters indicate significant differences among treatments. Note that the data presented in the figure do not statistically control for body mass (see text for statistical output).

**Table 2.** Model selection results for identifying the factors that explain variation in body composition of zebra finches. Only models with strong support (<2 ΔAIC_C_) are shown. Models were run as simple linear regressions and are ordered by ascending ΔAIC_C_.

| Models | K | AIC<sub>c</sub> | $\Delta AIC_c$ | $W_i$ | Acc $W_i$ | $R^2_{adj}$ | ER |
| --- | --- | --- | --- | --- | --- | --- | --- |
| **Carcass Fat Mass** |  |  |  |  |  |  |  |
| Temp+Mass+Sex | 5 | 131.74 | 0.00 | 0.21 | 0.21 | 0.205 | 1.00 |
| Sex*Mass+Temp | 6 | 131.98 | 0.24 | 0.19 | 0.40 | 0.226 | 1.13 |
| Temp*Mass+Sex | 6 | 132.75 | 1.02 | 0.13 | 0.52 | 0.213 | 1.66 |
| **Pectoral Mass** |  |  |  |  |  |  |  |
| Temp+Mass | 4 | 19.55 | 0.00 | 0.23 | 0.23 | 0.497 | 1.00 |
| Sex*Mass+Temp | 6 | 20.51 | 0.95 | 0.15 | 0.38 | 0.518 | 1.61 |
| Temp+Mass+Sex | 5 | 20.59 | 1.03 | 0.14 | 0.52 | 0.501 | 1.68 |
| Temp+Diet+Mass | 5 | 21.53 | 1.97 | 0.09 | 0.60 | 0.491 | 2.68 |
| **Visceral organ mass (PC1-Scores)** |  |  |  |  |  |  |  |
| Temp+Diet+Mass | 5 | 135.06 | 0.00 | 0.25 | 0.25 | 0.558 | 1.00 |
| Temp+Diet+Mass+Sex | 6 | 136.24 | 1.18 | 0.14 | 0.39 | 0.562 | 1.81 |
| Temp+Mass | 4 | 137.03 | 1.96 | 0.09 | 0.48 | 0.520 | 2.67 |

Note: Visceral organs (kidney, liver, heart, small intestine, and gizzard) were collapsed into a single axis (PC1) using a principal component analysis. Adjusted $R^2 = R^2_{adj}$ , “Mass” refers to body mass; all other abbreviations are as in Table 1.

### Pectoralis and visceral organ mass varied with acclimation temperature but not diet

Mass of the pectoralis was best predicted by temperature and body mass, which appeared in all models with strong support (ΔAIC_C_ < 2.0; Table 2). Despite diet and sex appearing in models with strong support (Table 2), neither term significantly predicted pectoralis muscle mass (both P >0.60, Table A2). Individuals acclimated to cool conditions (SFA-Cool and UFA-Cool) had a significantly heavier pectoralis than individuals acclimated to thermoneutrality (Fig 2B; Temperature: β = -0.267, P=0.002, Table A2). The pectoralis significantly increased with increasing body mass (Mass: β = 0.117, P<0.001, Table A2).

Visceral organ mass (PC1-scores) was best predicted by temperature and body mass, which appeared in all models of strong support; diet appeared in two of the top three models (Table 2). Based on model averaging, birds acclimated to cool conditions had significantly heavier visceral organs than those acclimated to thermoneutral conditions (Fig 2C; β = -1.975, p<0.001, Table A2). Not surprisingly, visceral organ mass significantly increased with increasing body mass (β = 0.266 p=0.003, Table A2). Despite dietary fat appearing in the top two models with strong support (Table 2), there was no statistically significant difference in the organ masses of birds on the UFA and SFA diets (Fig. 2C; β = -0.587, P= 0.182, Table A2). Similarly, although sex appeared in the second top model (ΔAIC_C_ = 1.18, Table 2), organ mass did not differ significantly between males and females (β = -0.123, p = 0.660, Table A2).

### Basal metabolic rate (BMR) was highest among individuals acclimated to a cool temperature

Variation in BMR was best predicted by acclimation temperature, body mass, and diet (ΔAIC_C_<2.0; Table 3); however, the model that contained only temperature and body mass was 1.70 times more likely to be the top model than one containing diet (ER=1.70, Table 3). Model averaging showed that BMR significantly increased with increasing body mass (β = 0.017, P < 0.001, Table A3) and was significantly higher among birds acclimated to cool temperatures than those acclimated to the thermoneutrality (Fig 3; Temperature: β = -0.138, P <0.001, Table A2). Diet did not significantly impact BMR (β =0.022, P = 0.234, Table A3).

**Figure 3:**
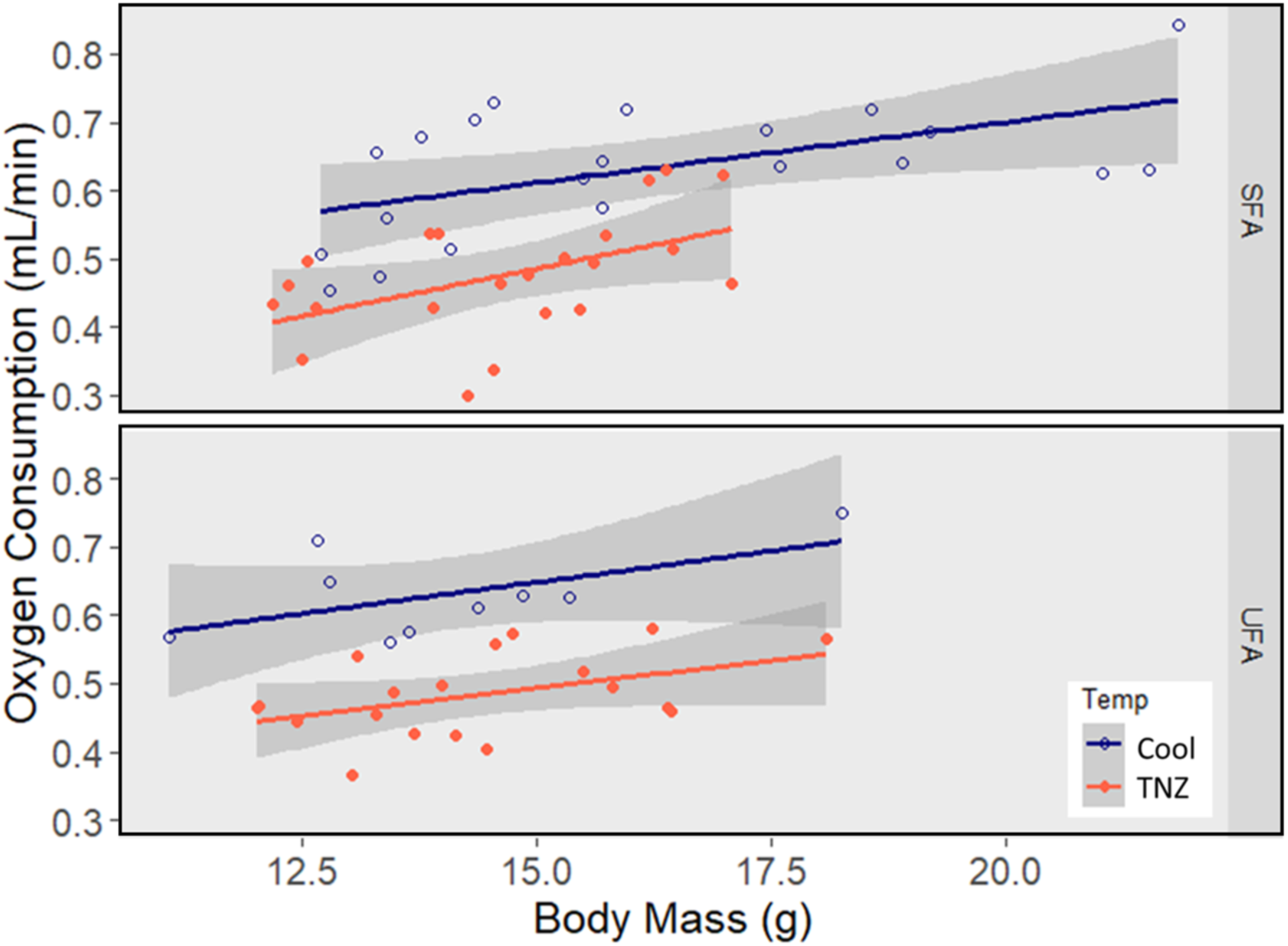
Basal metabolic rate (measured as rate of oxygen consumption) of zebra finches after a 3-months of acclimation to a diet enriched in saturated (SFA) or unsaturated fats (UFA), while living at thermoneutral (TNZ; 35 °C) or cool (24 °C) conditions. Birds living in the cool conditions had significantly higher BMR for a. given mass than those at thermoneutrality, irrespective of dietary treatment. 95% confidence intervals are depicted as grey areas around each treatment’s respective line of best fit.

**Table 3:**
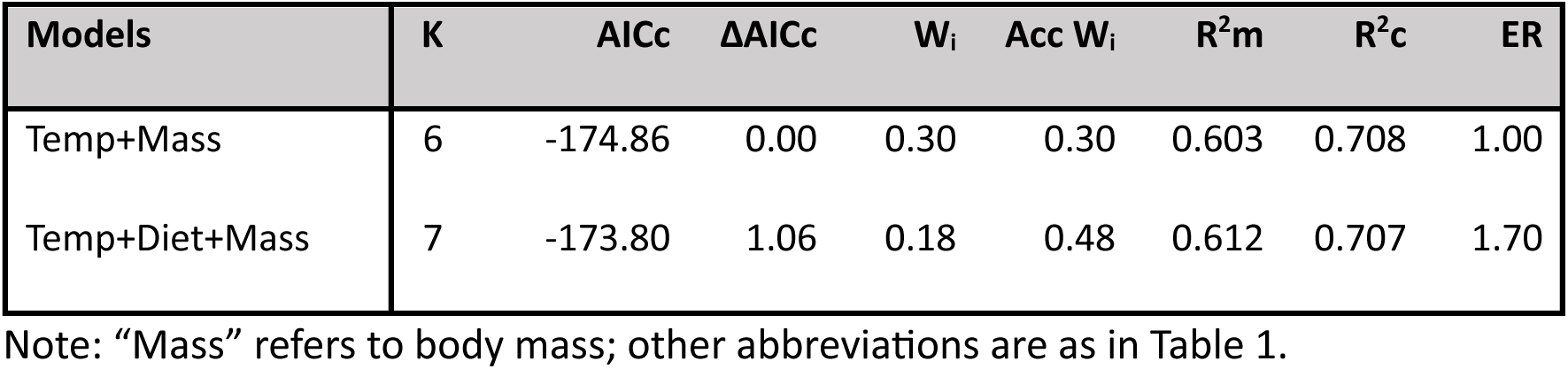
Model selection results for determining factors that explain variation in basal metabolic rate of zebra finches. Models with strong support (<2 ΔAIC_C_) are displayed. Models were run as linear mixed models using housing cage and respirometry chamber identity as random effects. Models were ordered by ascending ΔAIC_C_.

### Proportion of time spent feeding

Individuals spent a relatively low proportion of observation periods feeding (mean=5.6%, median=4.2%). There was no effect of temperature (β_TNZ_=0.027, n_cool_=23, n_TNZ_=40, p=0.634), diet (β_UFA_=-0.074, n_SFA_=37, n_UFA_=26 p=0.310), nor an interaction between temperature and diet (p=0.864) on proportion of time spent feeding (Table A4).

## Discussion

We studied the interactive effects of ambient temperature and diets differing in fatty acid content on the morphology and physiology of zebra finches over a chronic time frame. We hypothesized that the fatty acid content of the diet impacts the way individuals respond physiologically to a temperature change. We predicted that birds acclimated to cool temperatures would increase their body mass, presumably due to increased fat mass, flight muscles (pectoralis major) and visceral organs. Additionally, we predicted that birds exposed to cool temperatures and fed diets rich in unsaturated fatty acids would have smaller organs than those fed saturated fatty acids; we reasoned that incorporation of unsaturated fatty acids into cell membranes (and resultant heat generation due to leak) would reduce the necessity for organ hypertrophy. Accompanying predicted changes in visceral organ and pectoralis mass, we also predicted basal metabolic rate would be higher in the cool treatment, and higher in the UFA-fed birds. We found some support for our predictions; however, acclimation temperature had a far greater impact than diet over a chronic time frame.

It could be argued that any differences we detected were due to differences in the energy intake rates of the birds. However, we do not think this was the case. The sunflower oil and the coconut oil we used both contained 80 calories/10 mL of oil, and therefore the same calories between diet treatments. Further, we could detect no significant effect of diet, temperature, or their interaction on feeding frequency of individuals. As a result, we suggest our results reflect the impact of the differences in the composition of the fatty acids of the diet, rather than variation in energy intake rates.

### Body mass varied with temperature, diet, and length of acclimation

The body mass of individuals varied depending on the interactive effects of diet, the temperature at which they were housed, and the duration of the acclimation period. These interactive effects were most pronounced at 1-month, when individuals experiencing cool conditions lost mass if fed an unsaturated fatty acid rich diet (UFA-Cool), but not if fed a SFA-rich diet (SFA-Cool). In contrast, zebra finches housed at thermoneutrality maintained an intermediate mass, irrespective of diet.

Our data collected at 1-month contrast with those of Andersson et al (2018). In their study, wild caught great tits held at 3°C gained mass, irrespective of the fatty acid content of their diet. Typically, one of the initial responses exhibited by birds in response to cool temperature is an increase in fat mass, which relies on the dietary fats being consumed (West & Meng 1968; Carey et al. 1978). UFAs are easier to mobilize compared with equally long SFA carbon chains (Hulbert et al. 2005), but UFAs break down more readily due to lipid peroxidation (Zielinski & Pratt 2017). The molecular properties of UFAs may have caused UFA-cool zebra finches to have had more difficulty maintaining energy reserves, resulting in their initial mass loss despite ad lib food. In contrast, SFA-cool individuals maintained a relatively constant mass, with a (non-significant, p = 0.07) trend toward being heavier than all other groups after 6-months of exposure to cool temperatures.

Songbirds can acclimate to temperatures in relatively short periods of time (1-3 weeks; McKechnie 2008; Dubois et al. 2016; Thompson et al. 2016), including changes in mitochondrial activity and density (Zheng et al. 2013; Milbergue et al. 2022). In contrast, responding to changes in dietary fat composition may take months before birds fully acclimate their tissues to reflect the diet being consumed (Carter et al. 2019). Future studies could explore differences at the mitochondrial level in muscles or visceral organs following short-and long-term acclimation to diet and temperature, providing more information on specific physiological traits these birds developed to cope with chronic temperature change.

### Body composition was affected by temperature but not diet

When birds experience cold temperatures, they typically develop increased subcutaneous fat stores which act as both energy reserves and as a form of insulation (Houston et al. 1997; Dubois et al. 2016; Thompson et al. 2016). In our study, ambient temperature appeared in models with strong support for predicting variation in lipid mass (Table 2), however, based on model averaging, temperature was not a significant predictor (Table A2). Similarly, diet was not in any of the top models that explained variation in lipid mass. Why we failed to detect a chronic effect of either temperature or diet on stored lipid levels is not clear. However, most studies to date that link consumption of dietary fats and fat mass in birds are of migratory species and reflect short-term adjustments (Guglielmo et al. 2002; McWilliams et al. 2004; Maillet & Weber 2006; Dick et al. 2024). In preparation for migration, migratory species consume unsaturated fatty acids (especially polyunsaturated fatty acids) which are stored and then subsequently oxidized during migration (Pierce & McWilliams 2005; Maillet & Weber 2006). By rapidly exhausting their lipid stores, migratory species limit their exposure to UFAs and lower their risk of lipid peroxidation (Maillet & Weber 2006; Jensen et al. 2020; Lee et al. 2022). Our results may differ from previous studies because zebra finches are non-migratory, and we studied them over chronic rather than acute time frames. It would be interesting to look at the time course over which fat stores may change in response to temperature and diet using non-invasive monitoring (e.g., Guglielmo et al., 2011).

When zebra finches were exposed to cool conditions, they developed significantly heavier pectoral muscles and visceral organs than individuals held at thermoneutrality, irrespective of the fatty acid content of their diet (Fig 2B, 2C). Muscle hypertrophy is a common response in birds following cold acclimation, as the pectoralis is responsible for shivering thermogenesis (Vézina et al. 2006; Liknes & Swanson 2011) and acts as a thermal buffer to reduce heat loss (Houston et al. 1997; Thompson et al. 2016). Visceral organs have relatively high mass-specific-metabolic rates (Weber & Piersma 1996; Chappell et al. 1999; Wang et al. 2001), and an increase in size allows for greater heat production (Liknes & Swanson 2011; Zheng et al. 2013; Milbergue et al. 2018). We predicted there would be a statistical interaction between acclimation temperature and fatty acid content of the diet on muscle and visceral organ mass. Among individuals fed a UFA diet, we expected that UFAs would be incorporated into cell membranes, resulting in increased membrane leakiness (Hulbert et al. 2002; McWilliams et al. 2004; Maulucci et al. 2016) and heat release (Chaînier et al. 2000; Pani & Bal 2022). As a result, we expected that individuals acclimated to cool temperatures and fed UFAs would have smaller organs than individuals fed SFAs, as these individuals would be able to maintain their core temperature without the necessity of organ hypertrophy. However, this was not what we found, at least at the whole organ level, where effects of acclimation temperature outweighed any impacts of diet. It is possible that dietary fatty acids were simply not incorporated into the tissue membranes, however, this seems unlikely given previous studies (e.g. Ben-Hamo et al 2011). More likely is that the statistical effect of acclimation temperature and body mass on muscle and organ size was sufficiently large that it became difficult to detect any remaining statistical effect of diet.

### Basal metabolic rate was affected by temperature but not diet

Zebra finches acclimated to cool temperatures had higher BMR than those acclimated within their thermoneutral zone (when measured at 3-months), consistent with their larger pectoral muscles and visceral organs (at 6-months). Changes in BMR following chronic temperature acclimation has been widely reported (Liknes & Swanson 1996; McKechnie 2008; Dubois et al. 2016). However, BMR has also been shown to interact with dietary fat content during an “acute” time frame (Andersson et al. 2018). During exposure to a cold stress, great tits increased their BMR while fed SFA-rich diets relative to those fed an UFA-diet (Andersson et al. 2018). This was attributed to the birds on the SFA-diet birds converting SFAs into MUFAs, with an elevated BMR due to upregulation of enzymes associated with fatty acid biosynthesis and/or higher rates of oxidation of MUFAs. Nonetheless, over chronic time frames we detected no effect of diet, either in isolation or via an interaction with temperature. It is possible that the processes that contribute to an increase in BMR over acute time frames (e.g., upregulation of enzyme activity, Andersson et al 2018) play a relatively minor role over a chronic time frames.

## Conclusion

We demonstrated that over a chronic time frame zebra finches experiencing cool temperatures increased their basal metabolic rate, and adjusted their body composition, providing them with a long-term mechanism for maintaining core body temperature. Interestingly, the fatty acid content of their diet had no long-term effect on energy expenditure or body composition, at least at the level of biological organization we considered. Our study highlights that while it is important to understand how combined ecological factors may alter an individual’s physiology, it is also important to consider the duration over which acclimation occurs. Range shifts may cause species to experience long-lasting changes in food availability, either encountering novel food sources or losing major staples upon which they rely on to survive (Robb et al. 2008; Lu et al. 2012; Tekwa et al. 2022). Although our study provides insights into the long-term physiological consequences of differing diets and temperatures, future studies could incorporate a level of unpredictability. For example, daily fluctuations in weather, or changes in the demands and availability of food over seasonal time frames, may disrupt an individual’s capacity for acclimation (Wu et al. 2015). The consequences of such unpredictability on the capacity for acclimation remain largely unexplored.

## Acknowledgements

We thank Jason Allen and the staff in the Trent University Animal Care Facility for assistance with caring for the birds and Grant McClelland for discussion on fatty acids. Funding was provided by a Natural Sciences and Engineering Research Council (NSERC) – Collaborative Research and Training Grant and Research grant, and NSERC-Discovery Grants (awarded to GB and GM).

## Data Accessibility

All data will be archived on the Dryad Data repository

## Appendices

**Table A1:** Conditionally averaged body mass model estimates of predictor variables impacting changes in body mass using models with strong support (ΔAIC_C_ <2). All models were compared as linear mixed models using cage and individual identity (nested in cage identity) as random effects. Square brackets indicate the category being referred to for comparisons. Time values (1-month and 6-months) were all evaluated in reference to pre-experimental measurements (i.e., T_initial)_. Bolded p-values indicate statistical significance.

| Predictor Variables | Estimate | 95% CI | z value | Pr(>\|z\|) |
| --- | --- | --- | --- | --- |
| **(Intercept)** | 15.704 | 14.698, 16.711 | 30.572 | <0.001 |
| 1-month | -0.028 | -0.659, 0.603 | 0.087 | 0.9307 |
| 6-months | -0.485 | -1.293, 0.323 | 1.177 | 0.2392 |
| Temp [Warm] | -0.326 | -1.839, 1.186 | 0.423 | 0.6722 |
| Diet [UFA] | -0.702 | -2.185, 0.780 | 0.928 | 0.3532 |
| Diet:Temp | 0.137 | -2.202, 2.477 | 0.115 | 0.9086 |
| Temp:1-month | -0.823 | -1.728, 0.081 | 1.785 | 0.0743 |
| **Temp:6-months** | **-1.053** | **-2.047, -0.059** | **2.077** | **0.0378** |
| **Diet:1-month** | **-1.626** | **-2.634, -0.617** | **3.159** | **0.0016** |
| Diet:6-months | -1.084 | -2.665, 0.496 | 1.345 | 0.1787 |
| **Diet:Temp:1-month** | **2.512** | **1.158, 3.867** | **3.635** | **<0.001** |
| Diet:Temp:6-months | 1.494 | -0.358, 3.347 | 1.581 | 0.1139 |
| Sex [Male] | 0.330 | -0.886, 1.546 | 0.531 | 0.5952 |

**Table A2:** Model averaging estimates of predictor variables impacting changes in carcass fat, pectoral mass, and visceral organ mass (Vis PC1), using models with strong support (ΔAIC_C_<2). Dependent variables were examined as linear regressions. Square brackets indicate the category being referred to for comparisons. Bolded p-values denote statistical significance.

| Predictor Variables | Estimate | 95% CI | z value | Pr(>\|z\|) |
| --- | --- | --- | --- | --- |
| **Carcass fat mass** |  |  |  |  |
| (Intercept) | -0.524 | -3.583, 2.534 | 0.336 | 0.7369 |
| Temp [TNZ] | 0.578 | 0.092, 1.181 | 1.793 | 0.0730 |
| Mass | 0.150 | -0.149, 0.449 | 0.984 | 0.3249 |
| Sex [Male] | -0.766 | -4.804, 0.3271 | 0.372 | 0.7099 |
| Mass:Sex | 0.157 | -0.107, 0.712 | 0.735 | 0.4623 |
| **Pectoral muscle mass** |  |  |  |  |
| (Intercept) | 0.829 | -0.081, 1.738 | 1.785 | 0.0742 |
| **Temp [TNZ]** | **-0.267** | **-0.436, -0.097** | **3.080** | **0.0021** |
| **Mass** | **0.117** | **0.050, 0.185** | **3.413** | **0.0006** |
| Sex [Male] | -0.276 | -1.886, 0.711 | 0.511 | 0.6094 |
| Mass:Sex | 0.018 | -0.021, 0.169 | 0.450 | 0.6525 |
| Diet [UFA] | -0.009 | -0.230, 0.111 | 0.219 | 0.8263 |
| **Visceral organ mass (PC1-scores)** |  |  |  |  |
| (Intercept) | -2.071 | -4.768, 0.627 | 1.505 | 0.1324 |
| **Temp [TNZ]** | **-1.975** | **-2.752, -1.199** | **4.985** | **<0.001** |
| Diet [UFA] | -0.587 | -1.453, -0.004 | 1.336 | 0.1816 |
| **Mass** | **0.266** | **0.089, 0.443** | **2.946** | **0.0032** |
| Sex [Male] | -0.123 | -1.165, 0.307 | 0.440 | 0.6596 |

**Table A3:** Conditionally averaged model estimates of predictor variables impacting BMR using models with strong support (ΔAICC <2). Models were examined as linear mixed models using cage identity and respirometry chamber as random effects. Square brackets indicate the category being referred to for comparisons. Bolded p-values denote statistical significance.

| Predictor Variables | Estimate | 95% CI | z value | Pr(>\|z\|) |
| --- | --- | --- | --- | --- |
| (Intercept) | **0.372** | **0.251, 0.492** | **6.058** | **<0.001** |
| Temp [TNZ] | **-0.138** | **-0.175, -0.100** | **7.114** | **<0.001** |
| Mass | **0.017** | **0.010, 0.024** | **4.592** | **<0.001** |
| Diet [UFA] | 0.022 | -0.014, 0.059 | 1.190 | 0.2341 |

**Table A4:** Results of a linear mixed model for the proportion of observations where birds were observed feeding. Data were arcsine square root transformed prior to analysis. A single individual with an unusually high proportion of time spent feeing was removed as a possible outlier (>4SD above the mean). Cage identity was included as a random effect. Reference treatments are in brackets. (n_SFAcool_=15, n_SFATNZ_=22, n_UFAcool_=8, n_UFATNZ=_18)

| Predictor Variables | Estimate | CI | T value | Pr(>\|z\|) |
| --- | --- | --- | --- | --- |
| **(Intercept)** | **0.227** | **0.155, 0.297** | **5.632** | **<0.001** |
| Temp (TNZ) | 0.027 | -0.070, 0.125 | 0.492 | 0.634 |
| Diet (UFA) | -0.074 | -0.195, 0.049 | -1.069 | 0.310 |
| Temp:Diet | 0.015 | -0.140, 0.170 | 0.176 | 0.864 |

